# Peer presence and familiarity as key factors to reduce cocaine intake: an effect mediated by the Subthalamic Nucleus

**DOI:** 10.1101/2021.06.08.447497

**Authors:** E Giorla, S Nordmann, C Vielle, Y Pelloux, P Roux, C Protopopescu, C Manrique, K Davranche, C Montanari, L Giorgi, A Vilotitch, P Huguet, P Carrieri, C Baunez

## Abstract

Stimulant use, including cocaine, often occurs in a social context whose influence is important to understand to decrease intake and reduce associated harms. Given the regulatory role of the subthalamic nucleus (STN) on cocaine intake and emotions, we investigate its role on such influence of social context on cocaine intake. We explored the influence of peer presence and familiarity on the frequency of self-administered cocaine and its neurobiological basis. We first compared cocaine intake in various conditions (alone or with peers with different characteristics: observing or self-administering, familiar or not, cocaine-naive or not, dominant or subordinate) in rats (n=90). The risk of drug consumption was reduced when a peer was present, observing or self-administering as well, and further diminished when the peer was unfamiliar (vs familiar). The presence of a cocaine-naive peer further decreased cocaine consumption. The presence of a non-familiar and drug-naive peer represents thus key conditions to diminish cocaine intake. We tested the effects of STN lesions in these various conditions and also conducted social experiments to validate the role of STN in social cognition. The STN lesion by itself reduced cocaine intake to the level reached in presence of a stranger naïve peer and affected social cognition, positioning the STN as one neurobiological substrate of social influence on drug intake. Finally, with a translational research approach, we compared the drug intake in these conditions in human drug users (n=77). This human study confirmed the beneficial effect of social presence, especially of strangers. Our results indirectly support the use of social interventions and harm reduction strategies, in particular supervised consumption rooms for stimulant users.

## Introduction

Recreational drug intake often takes place in a social environment. The social context is known to influence initiation^1,2^, persistence^3,4^, increase^5^ and cessation^6,7^ of drug consumption^8,9^. It is also a major determinant of risk practices, such as sharing injecting equipment^10^. A social environment encompasses two types of social factors: distal social factors (i.e. the consumer’s broader social environment, which is not present immediately during drug consumption) and the proximal social factors (i.e. the immediate social environment of the consumer, during the drug-taking).

Like humans, rats live in complex social environment^11^ showing a large panel of social abilities and behaviors^12^, and offering suitable models to elucidate neural processing of drug intake. Interestingly, the influence of distal social factors has been widely studied and, in rats, they mirror the effects highlighted in humans. Likewise, in these two species, stress, isolation and rejection are associated with higher rates of drug use^13–18^. In contrast, strong familial ties in humans^19–22^, and enriched environment in rats, are associated with lower rates of drug^15–17,23,24^.

Less is known about the proximal social influence on drug use, and studies on rodents may have contradicting results. Indeed, social interactions may act as an alternative reinforcement over drug use, diverting animals from consumption^25–27^, but they can also have a facilitating effect on drug consumption^28–31^. Such discrepancy between results may originate from differences in experimental conditions. Notably, the presence of a peer during alcohol consumption has been shown to differently modulate rats’ consumption, depending on the familiarity status of the present peer^32^. Similarly, the broader drug experience of this peer present during self-administration could differently affect the consumption^33^. However, in these animal studies, many characteristics of the peers such as familiarity, dominance status, former experience of the drug, were not all investigated systematically, nor the neurobiological mechanisms and brain structures involved in the influence of proximal social factors on addiction.

Recent studies on the neuroanatomical basis of the social modulation of drug use have focused on the reward system. Among them, the Nucleus Accumbens (NAc), the amygdala and the insular cortex are involved in both drug and social interaction reinforcing effects (for review see Pelloux et al 2019^34^). The Subthalamic Nucleus (STN), deep cerebral structure belonging to the basal ganglia^35^, has opposite effects on food and drug motivation^36^ and is implicated in social emotional processing^37^. Thus, it could play a key role in the influence of social proximal factors on drug consumption. Moreover, in former studies, it has been shown that STN lesion impairs the expression of both positive and negative emotional state^38^ and abolishes the influence of positive and negative ultrasonic vocalizations on cocaine intake^39^, bringing some clues regarding STN contribution to emotional processing. Since we assumed that the social context and the effect of the presence of a peer on drug intake implies brain structures involved in emotional processes, we have thus focused on the role of STN the influence of social context on drug consumption and in social interactions.

Epidemiologic studies have also shown an impact of a peer presence on alcohol consumption^41^ and an increased risk for a teenager of becoming a drug user if their friends also consume drugs^42^. But, to our knowledge, no study to date has specifically focused on the influence of peer characteristics and drug-using status and their effect on cocaine consumption in humans. The few existing studies in this area only examined the influence of peer presence and close relationships on outcomes such as alcohol use and craving during stressful events^43^. To validate this model, there is a need for a translational approach, confirming if the same processes are involved, in both humans and rats, regarding the proximal social influence on drug consumption.

In this study, we aimed thus to characterize how the dyad characteristics (familiarity, shared history of drug exposure) could influence cocaine intake and how it could be modulated by STN lesions in self-administering rats. We also investigated the role of STN on social preference to understand how social interaction could be rewarding for adult rats and therefore influence drug consumption. Finally, with a translational research approach, we ran an epidemiologic study in human drug users, allowing us to control that the experimental conditions used for rats mirrored the processes observed in humans.

## Results

### Experiment 1: Influence of the presence of peer and its characteristics on cocaine self-administration in sham and STN lesioned rats

#### Influence of peer presence

##### Observing peer

Sham control rats in the “alone” condition took an average of 13.51 (± standard error of the mean (SEM: 0.82) cocaine injections during the one-hour self-administration sessions. When a peer was present (familiar or not), the average value decreased to 10.51 (±0.72) (**Figure 2A**). This translates to a 22% decrease in the relative risk of cocaine intake when rats were in the presence of an observing peer, compared with when they were alone (IRR [95% CI]=0.78 [0.71-0.86], p<10^−3^) (see supplemental **Table S2A**). When the STN lesioned rats were alone in the self-administration chamber, they took an average of 12.07 (±0.59) injections. There was thus no significant difference between the sham and the STN lesioned groups in the “alone” condition (p=0.527, group effect, mixed effects Poisson regression). When an observing peer was placed in the chamber separated by a grid, the STN lesioned rats took less cocaine injections than when alone (means 6.63 (±0.53)) (**Figure 2C**). This translates to a 45% decrease in the relative risk of cocaine intake when rats were in the presence of an observing peer, compared with when they were alone (IRR [95% CI]=0.55 [0.50-0.61], p<10^−3^) (see supplemental **Table S2A**).

**Figure 1.**
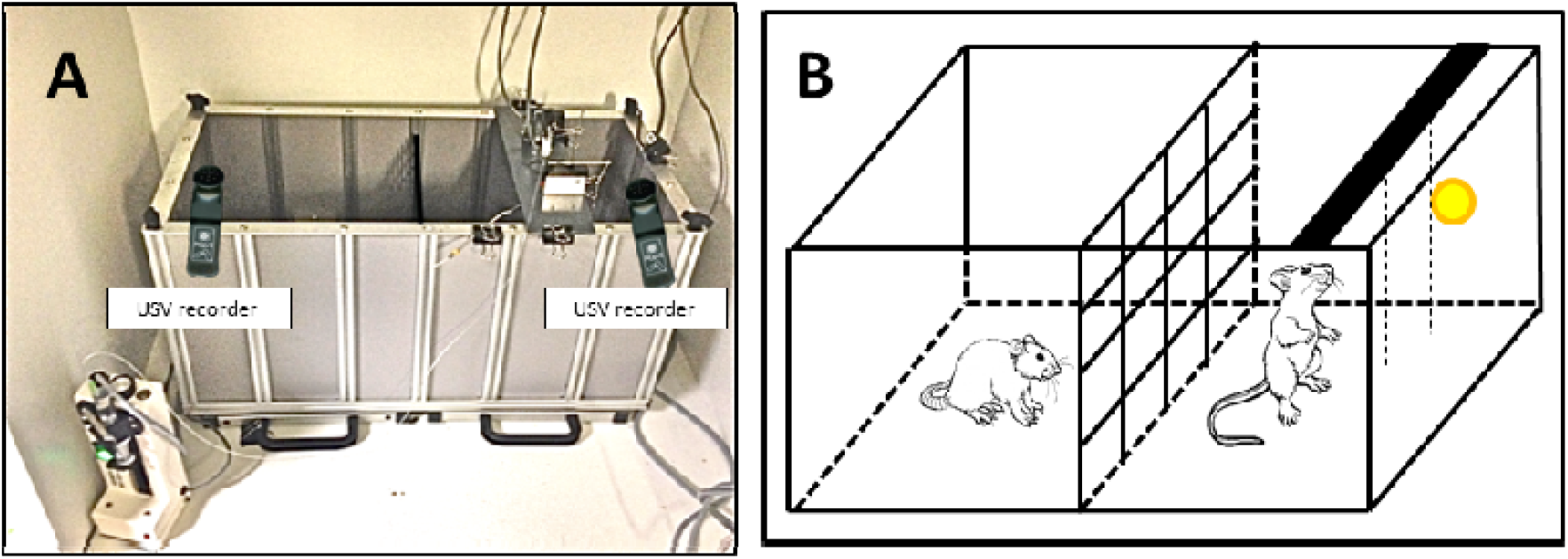
Picture and schematic representation of the self-administration experimental apparatus. **A**. Picture of one experimental box from above. **B**. Schematic diagram of the same experimental box from the side. The observing rat is in the left compartment separated by a grid (2,5×2,5cm, allowing visual, auditive, olfactory and limited tactile interactions between the 2 rats), while the self-administering rat is in the right compartment and has access to two chains (small dotted lines), one of which is the active chain and leads to cocaine delivery.

**Figure 2.**
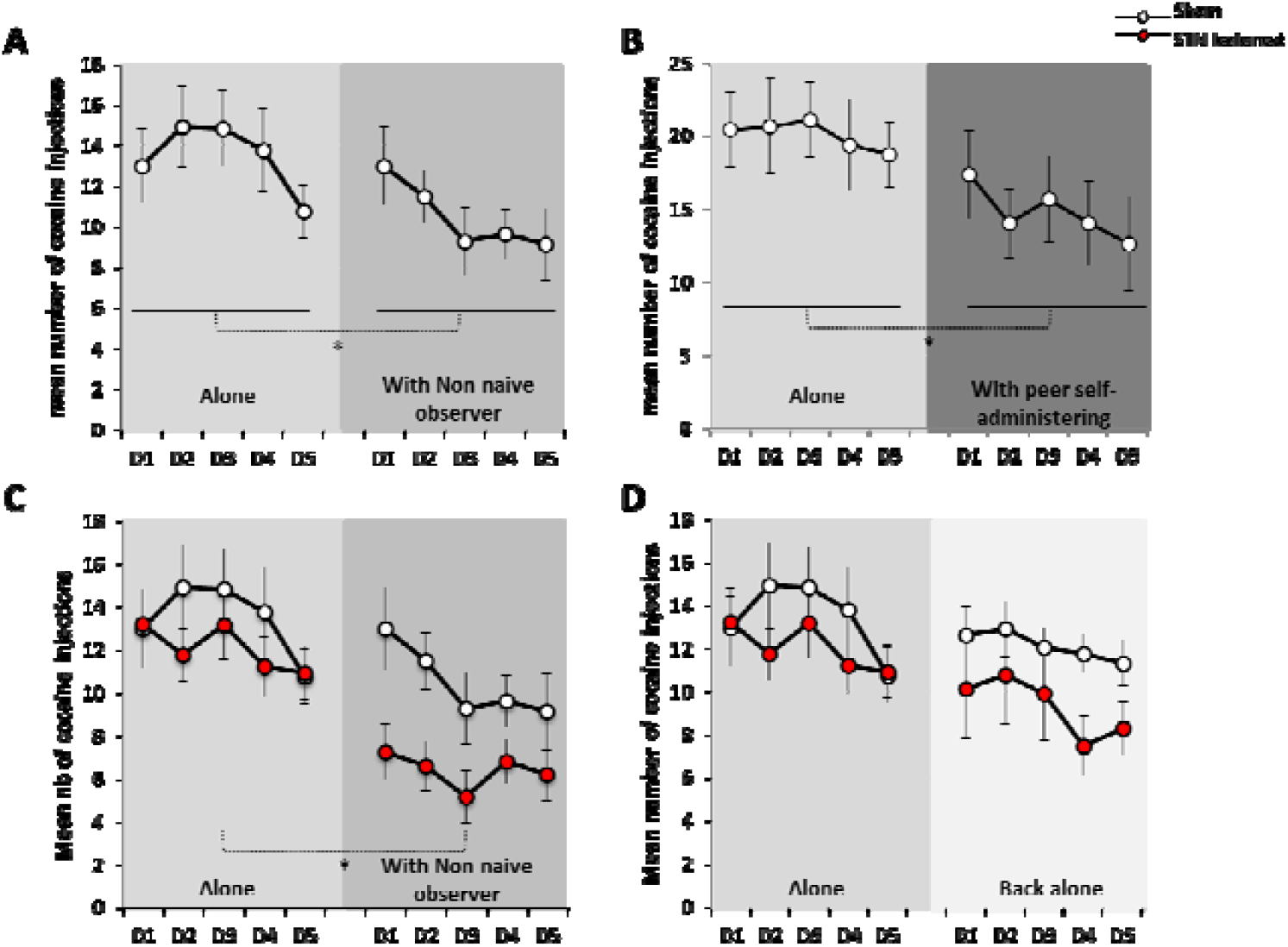
Influence of the presence of an observing or self-administering peer on cocaine consumption in sham and STN lesioned rats. **A. Influence of the presence of an observing peer on cocaine self-administration in rats**. The results are illustrated as the mean (± SEM) number of cocaine injections (80µg/90µl/injection) per 1h-session during 5 consecutive sessions of baseline (“alone”, D1 to D5) and during 5 consecutive sessions in presence of a peer (“with non-naive observer”, D1 to D5), for sham control rats (white dots, n=14) (*p<0.05 group effect, Poisson regression). **B. Influence of the presence of a peer also self-administering cocaine on cocaine self-administration**. The results are illustrated as the mean (± SEM) number of cocaine injections (80µg/90µl/injection) per 1h-session during 5 consecutive sessions of baseline (“alone”, D1 to D5) and during 5 consecutive sessions in co-administration (“with peer self-administrating rat”, D1 to D5, n=14). (*p<0.05 group effect, Poisson regression). **C. Effect of the STN lesion on the influence of the presence of an observing peer on cocaine self-administration in rats**. The results are illustrated as the mean (± SEM) number of cocaine injections (80µg/90µl/injection) per 1h-session during 5 consecutive sessions of baseline (“alone”, D1 to D5) and during 5 consecutive sessions in presence of a peer (“with non-naive observer”, D1 to D5), for sham control rats (white dots, n=14) and for the STN lesioned rats (red dots, n=17), (*p<0.05 group effect, Poisson regression). **D. Return to baseline cocaine consumption when rats are tested alone again**. The results are illustrated as the mean (± SEM) number of cocaine injections (80µg/90µl/injection) per 1h-session for 5 consecutive days at baseline (“Alone”, D1 to D5: day 1 to 5 sham n=14 STN lesioned n=17) and for 5 consecutive sessions alone after the experience of being in the presence of a non-naive peer (“Back alone”, D1 to D5: day 1 to 5 sham n=14 STN lesioned n=17).

The reduced cocaine intake induced by the presence of a peer was stronger in STN lesioned rats (6.63 (± 0.53)) than in sham rats (10.51 (±0.72)) (p=0.016, group effect, mixed effects Poisson regression) (**Figure 2C**).

##### Self-administering peer

When the peer also had access to cocaine, we observed a comparable effect to the first part of the experiment where the present peer was only observing, with no access to the drug (**Figure 2B**). First, rats were alone to acquire the self-administration behavior and reached a stable number of cocaine injections (20.04 (±1.21)). When rats were in a presence of a peer also self-administering, they took an average of 14.84 injections (±1.27) (**Figure 2B**). This represents a 26% decreased risk of consuming cocaine when they were placed in presence of a peer also taking cocaine (IRR [95% CI]=0.74 [0.68-0.80], p<10^−3^) (see supplemental **Table S2B**) compared with when they were alone (see **Figure 5A** for levels of consumption).

Chronologically, in this study, after exposure to a non-naive (non-taking cocaine) peer, rats were tested again alone to self-administer (“back alone” condition) to be then tested in presence of a non-familiar naive of cocaine rat. In sham rats, when the animals were tested in the “back alone” condition after the test of social presence, cocaine consumption returned to a level close to the baseline (“alone”) level (11.87 (±0.48) versus 13.51 (±0.82)respectively) (**Figure 2D**).

#### Influence of peer familiarity

##### Observing peer

In sham controls animals, the average number of cocaine injections self-administered during the one-hour sessions was lower when the peer was “non-familiar” (7.60 (±0.89)) than when “familiar” (12.7 (±0.94)) (**Figure 3A**). The Poisson regression analysis showed a 15% decreased relative risk between alone and presence of a familiar peer (IRR [95% CI]=0.85 [0.76-0.96], p=0.009) (see supplemental **Table S2A**). This decrease was further significant (35%) in presence of a non-familiar peer (IRR [95% CI]=0.65 [0.55-0.77], p<10^−3^) and also when compared to the condition of presence of a familiar peer (non-familiar vs familiar peer: IRR=0.55 [0.35-0.87], p<0.011) (see supplemental **Table S2A**).

**Figure 3.**
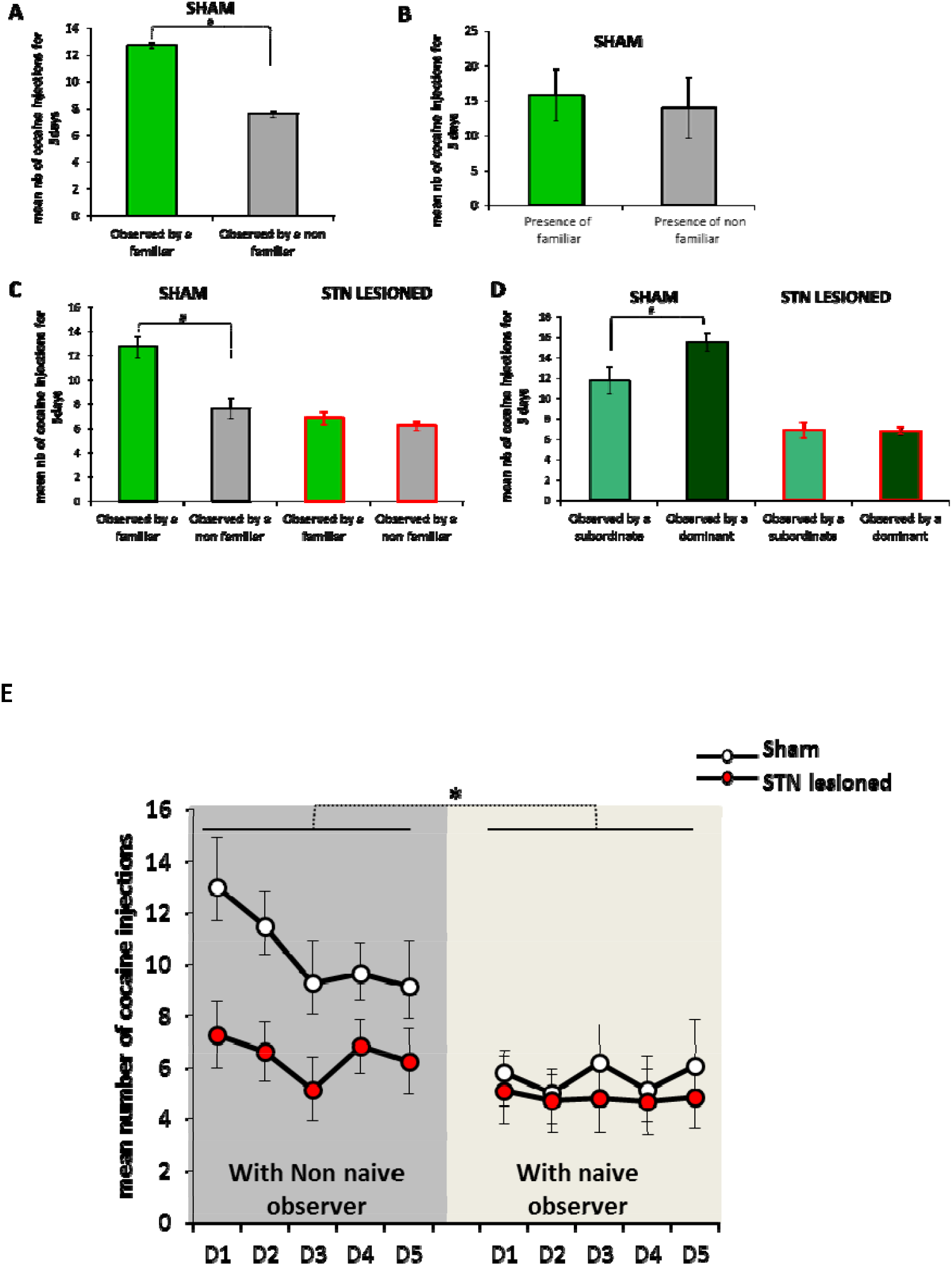
Influence of the familiarity, dominance status and history of drug exposure of a peer on cocaine consumption in sham and STN lesioned rats. A. **Influence of familiarity of the present peer on cocaine self-administration in rats**. The results are illustrated as the mean (± SEM) number of cocaine injections (80µg/90µl/injection) per session and averaged for 5 consecutive days of social interaction with either a familiar peer (green bar, sham n=8) or an **unknown** peer (“non-familiar”, grey bar, sham n=6), (#p<0.05 conditions effect, Poisson regression). **B. Influence of familiarity with a self-administering peer on cocaine self-administration in rats**. The results are illustrated as the mean (± SEM) number of cocaine injections (80µg/90µl/injection) per session and averaged for 5 consecutive days of self-administration when the rat was in presence of a peer also self-administering cocaine (self-administering partner was “familiar” green bar, n=6) or a stranger (“non-familiar”, grey bar, n=8)) **C. Effect of STN lesion on the influence of the familiarity of the present peer on cocaine self-administration in rats**. The results are illustrated as the mean (± SEM) number of cocaine injections (80µg/90µl/injection) per session and averaged for 5 consecutive days of social interaction with either a familiar peer (green bars, dark line sham n=8, red line STN lesioned n=6) or an **unknown** peer (“non-familiar”, grey bars, dark line sham n=6, red line STN lesioned n=11), (#p<0.05 conditions effects, Poisson regression). **D. Effect of the STN lesion on the influence of dominance status within familiar peers on cocaine self-administration in rats**. The results are illustrated as the mean (± SEM) number of cocaine injections (80µg/90µl/injection) per session and averaged for 5 consecutive days of social interaction with either its familiar subordinated peer (light green bars, dark line sham n=4, red line STN lesioned n=3) or its familiar dominant peer (dark green bars, dark line sham n=4, red line STN lesioned n=3), (#p<0.05 conditions effects, Poisson regression). **E. Influence of the peer history of drug exposure on cocaine consumption in sham and STN lesioned rats**. The results are illustrated as the mean (± SEM) number of cocaine injections (80µg/90µl/injection) per 1h-session for 5 consecutive sessions under observation of a non**-**familiar peer with a history of cocaine self-administration (left, “non **-**naive peer”, D1 to 5: day 1 to 5, white dots sham n=14; red dots STN lesioned n=17) and under observation of a non**-**familiar peer **naive** to cocaine (right, “**naive** peer”, D1 to D5: day 1 to 5, white dots sham n=11; red dots STN lesioned n=15); (*p<0.05 conditions effects, Poisson regression).

In the STN lesioned rats, no matter the characteristics of the peer, the cocaine intake decreased in the same manner to a low level (observed by familiar: mean 6.87 (±0.73), observed by a stranger: mean 6.51 (±0.73)) (**Figure 3C**). The Poisson regression analysis showed a 33% decreased relative risk between alone and presence of a familiar peer (IRR [95% CI]=0.57 [0.48-0.68], p<10^−3^) (see supplemental **Table S2A**). The reduction of risk to take cocaine was equivalent (36%) when the peer was non-familiar (IRR [95% CI]=0.54 [0.47-0.61], p<10^−3^), and no significant difference was found for the comparison between presence of an observing non familiar vs familiar peer (IRR=0.81 [0.44-1.51] p=0.517) (see supplemental **Table S2A**).

##### Self-administering peer

These results were also confirmed when rats self-administered cocaine in the presence of a peer which could also take cocaine (**Figure 3B**). In sham-control animals, the average number of cocaine injections self-administered during the one-hour sessions was lower when the peer was “non-familiar” (14.15 (±1.87)) than when “familiar” (15.79 (±1.60)) (**Figure 3B**). In the mixed Poisson model analysis, we found that compared with being alone, a decreasing relative risk of consumption was observed from being with a familiar peer (22% reduction) (IRR [95% CI]=0.78 [0.69-0.88], p<10^−3^) to being with a non-familiar peer (29% reduction)(IRR [95% CI]=0.71 [0.64-0.79], p<10^−3^) (see supplemental **Table S2B**). However, these decreases were smaller than those observed in the former experiment with observing peers.

#### Influence of the dominance/subordination relationship

Further analysis assessing the dominance status revealed that, in sham control animals, the average number of cocaine injections self-administered during the one-hour sessions was lower when the observing peer was the familiar-subordinate (11.74 (±1.15)) than when it was the dominant one (15.51 (±0.69)) (p=0.036, group effect, mixed effects Poisson regression) (**Figure 3D**). In STN lesioned rats, no significant difference was found when comparing the frequency of cocaine consumption in rats in the presence of the subordinate (6.86 (+-0.67)) and in the presence of the dominant (6.8 (+-0.69) familiar peer **(Figure 3D)**.

#### Influence of peer history of drug exposure

In the presence of a cocaine-naive peer, sham rats significantly reduced their drug intake to an average of 5.64 (±0.56) cocaine injections, in comparison with an average of 10.51 (±0.72) injections in the presence of a non-naive of cocaine peer (familiar to the subject and non-familiar together) (**Figure 3E**).

The Poisson model showed a progressive decrease in the risk of cocaine consumption from alone to the presence of a non-naive peer (22% reduction; IRR [95% CI]=0.78 [0.71-0.86], p<10^−3^) and from back alone to the presence of a naive peer (53% reduction; IRR [95% CI]=0.47 [0.41-0.54], p<10^−3^) (see supplemental **Table S2A**).

In STN lesioned rats, when tested in the “back alone” condition after the test of social presence, the return to baseline level was less obvious (mean 9.08 (±0.90) vs 12.07 (±0.59)) (**Figure 2D**). STN lesioned rats also significantly decreased their cocaine intake in presence of a cocaine naive peer (4.96 (±0.55)) when compared to presence of a non-naive peer (6.63 (±0.53)) (p<0,05, conditions effects, Poisson regression) (**Figure 3E**). The Poisson model showed a similar decrease in the risk of cocaine consumption from “alone” to the “presence of a non-naive peer” (45% reduction; IRR [95% CI]=0.55 [0.50-0.61], p<10^−3^) and from “back alone” to the “presence of a naive peer” (46% reduction; IRR [95% CI]=0.54 [0.48-0.61], p<10^−3^) (see supplemental **Table S2A**).

### Experiment 2: Reinforcing properties of Social presence

#### Social preference

To evaluate the potential reinforcing properties of social presence, we first tested if the presence of a peer was preferred over an object and whether the STN lesion could affect this discrimination. We thus subjected sham-control and STN-lesioned rats to a social preference test, using a social stimulus (a stranger rat or the cage-mate) and an object. As illustrated in **Figure 4A**, when the social stimulus was a stranger rat, both sham-control and STN-lesioned rats spent more time exploring the cage containing the rat than the one containing an object (sham n=7, W=28, p=0.0156 ; lesioned n=9, W=45, p=0.0039 (Wilcoxon matched-pairs signed rank test)). However, when the cage-mate was used as the social stimulus, only STN-lesioned rats showed a preferential exploration of the cage containing the social stimulus (W=45, p=0.0039 (Wilcoxon matched-pairs signed rank test) (**Figure 4B)**; sham-control group: W=20, p=0.1094 (Wilcoxon matched-pairs signed rank test)). Together, those results highlight the fact that a stranger rat is preferred over an object and that STN lesions seem to impair the dissociation between familiar and stranger peers.

**Figure 4.**
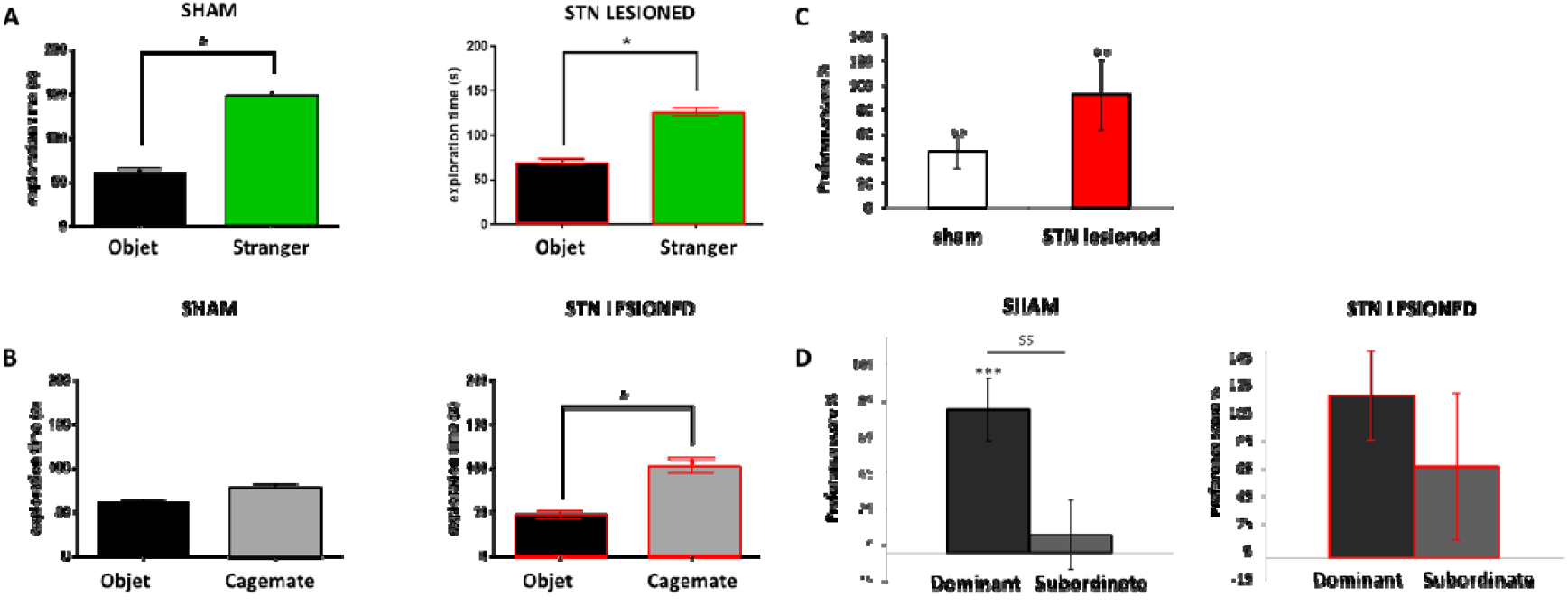
Effect of the STN lesion on the rewarding properties of social interactions. **A. Effect of the STN lesion on the social preference over an object: preference between a stranger peer over an object**. The mean exploration time (sec. (±SEM) i.e. the time spent exploring the social stimulus (green bar) and the object (black bar), is represented for sham rats (dark lines, n=7) and STN lesioned rats (red lines, n=9), (*: p<0.05 compared to stranger stimulus (Wilcoxon)) **B. Effect of the STN lesion on the social preference over an object: preference between a familiar peer over an object**. The mean exploration time (sec., (±SEM) i.e. the time spent exploring the social stimulus (grey bar) and the object (black bar), is represented for sham rats (dark lines, n=7) and STN lesioned rats (red lines, n=9), (*: p<0.05 compared to familiar stimulus (Wilcoxon)) **C. Conditioned place preference test for peer presence: effect of the STN lesion on the preference score for the presence of a peer**. The mean preference score (±SEM), i.e. the time spent in the compartment paired with the reinforcer on the testing day minus the time spent in this same compartment at day 1 before conditioning is represented for the sham control group (white bar, n=66) and for the STN lesioned group (red bar, n=14). ** p<0.001 compared to a theoretical zero (sign test) (p=0.082 group comparison Mann-Whitney sham vs STN lesioned). **D. Conditioned place preference test for peer presence: effect of the dominance status on the preference score for the presence of a peer for the control and STN lesion group**. The mean preference score (±SEM), i.e. the time spent in the compartment paired with the reinforcer on the testing day minus the time spent in this same compartment at day 1 before conditioning is represented for the dominant rats (black bars, dark line: sham n=33 ; red line: STN lesioned n=7,) and for the subordinate (dark grey bars, dark line: sham n=33; red line: STN lesioned n=7). *** p<0.001, ** p<0.01 compared to a theoretical zero (sign test); $$ p<0.01 significant group comparison (Mann Whitney dominant vs subordinate).

#### Conditioned place preference

To check whether or not, the presence of a peer is a positive social context with reinforcing properties, the conditioned place-preference paradigm was used with the presence of the cage-mate. In this paradigm, the rewarding effects of a given reinforcer (here, social interaction) are inferred by comparing the time spent in a specific environment previously paired with the reinforcer with the time spent in the same compartment before its association with the supposed reward (i.e. preference score).

First, we found no significant differences between sham control group and STN lesioned group on the preference score (group effect: p=0.082 (Mann-Whitney Test), **Figure 4C**). Nevertheless, we found a conditioning effect for both groups. Indeed, the preference score for these two groups was different from zero (conditioning effect: p<0.001, (sign test), **Figure 4C**). This suggests that adult sham control and STN lesioned rats significantly prefer the compartment associated with the presence of a conspecific.

Further analysis assessing the dominance status revealed that, in the sham control group, the preference score was only significant for the dominant rats (conditioning effect: p<0.0001 (sign test); dominance effect: p<0.001 (Mann-Whitney test), **Figure 4D**). Subordinate control rats did not develop any preference or aversion for the compartment associated with the presence of their dominant peer (conditioning effect: p=0.68, (sign test)).

Possibly due to inter-individual variability, in the STN lesioned group, dominant and subordinate rats did not show any significant differences in their preference score (dominance effect: p=0.186, (Mann-Whitney test), **Figure 4D**). Nevertheless, only dominant STN rats showed a preference score significantly different from zero (conditioning effect: p<0.05, (sign test), **Figure 4D**).

### Experiment 3: Translational towards human drug users

In humans, among episodes involving one other peer, we note that the latter was always a drug user (for episodes with a group, peers were not characterized). Whether the drug user was actually consuming or not during the same episode was not directly documented, but this is why it was important to perform a rat study in which the peer could also have access to the drug during the same episode.

In the human study, results showed a 37% decreased relative risk of cocaine intake during an episode when one peer was present with respect to alone (IRR [95% CI]=0.63 [0.42-0.94], p=0.023), after adjusting for unstable housing and other drug intake (stimulant, other including alcohol) (**Table 1** and **Figure 5A**). These results are in line with those from the rat study, where a decrease in relative risk of cocaine intake was observed in a presence of an observing peer (**Figure 5A**).

**Table 1.** **Human study** - Association between peer presence and cocaine use frequency among humans (N=77, 246 episodes of cocaine use) – multivariable analysis.

|  | Coefficient Estimate<br>[95% CI]* | IRR [95% CI]* | P-value |
| --- | --- | --- | --- |
| **Social proximal context** |  |  |  |
| Alone | 0 | 1 |  |
| With one peer | -0.469 [-0.873 ; -0.065] | 0.63 [0.42 ; 0.94] | 0.023 |
| Group | 0.119 [-0.246 ; 0.485] | 1.13 [0.78 ; 1.62] | 0.523 |
| **Unstable housing** |  |  |  |
| No | 0 | 1 |  |
| Yes | 0.822 [0.466 ; 1.178] | 2.28 [1.59 ; 3.25] | <10<sup>-3</sup> |
| **Daily stimulant intake** |  |  |  |
| No | 0 | 1 |  |
| Yes | 0.429 [0.129 ; 0.730] | 1.54 [1.14 ; 2.07] | 0.005 |
| **Other drug intake (alcohol included)** | 0.150 [-0.002 ; 0.302] | 1.16 [1.00 ; 1.35] | 0.053 |

**Figure 5.**
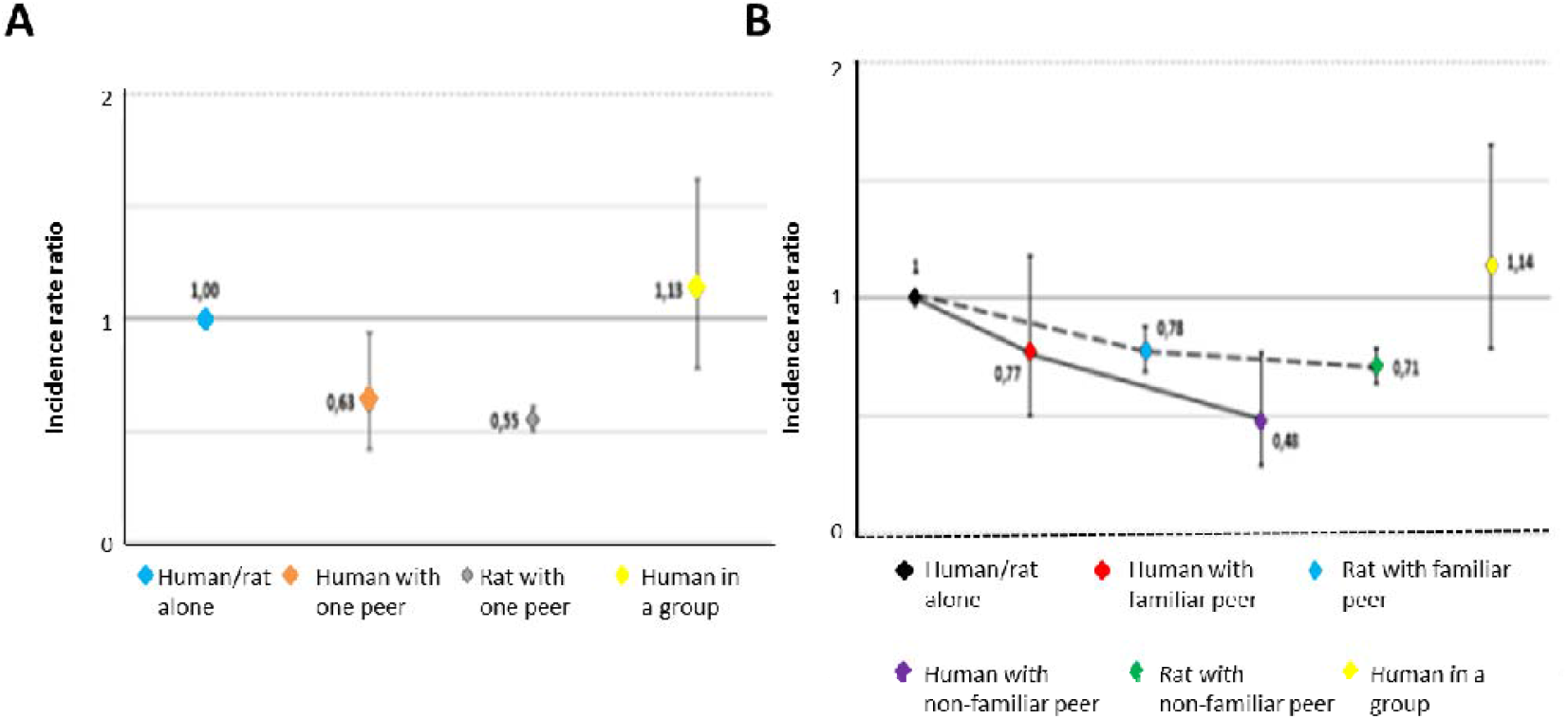
Translation towards human drug users. **A. Adjusted incidence rate ratios (relative risk) of frequency of drug consumption depending on the peer presence in humans and in rats** using the condition **“alone” as reference**. Blue, orange and yellow diamonds represent the adjusted incidence rate ratio from the multivariable analysis using GEE Poisson model in humans of the variable peer presence (reference=alone): human alone (blue diamond) and human with one peer (orange diamond) and group (yellow diamond). The lower and upper dashes represent, respectively, the lower and upper bounds of the 95% confidence interval. Blue and grey diamond represent the adjusted incidence rate ratio from the multivariable analysis using GEE Poisson model in rats of the variable peer presence (reference=alone): rat with one peer (grey diamond). The lower and upper dashes represent, respectively, the lower and upper bounds of the 95% confidence interval. **B. Adjusted incidence rate ratios (i.e. relative risk) for drug consumption depending on peer familiarity in humans and rats using the condition “alone” as reference**. Blue, orange and yellow diamonds represent the adjusted incidence rate ratio from the multivariable analysis using GEE Poisson model in humans of the variable familiarity (reference=alone; black diamond: human alone, red diamond: human with familiar peer, purple diamond: human with non-familiar peer). The lower and upper dashes represent, respectively, the lower and upper bounds of the confidence interval. Black, blue and green diamonds represent the adjusted incidence rate ratio from the multivariable analysis using GEE Poisson model in rats of the variable familiarity (reference=alone; blue diamond: rat with familiar peer, green diamond: rat with non-familiar peer). The lower and upper dashes represent, respectively, the lower and upper bounds of the confidence interval.

However, there was no significant difference in cocaine intake when a group of peers was present compared with being alone (p=0.523) (**Table 1**).

As for the rat experiment, we also observed a decrease in relative risk of cocaine use during an episode from familiar peer presence (IRR [95% CI]=0.77 [0.50-1.18], p=0.233) to non-familiar peer presence (IRR [95% CI]=0.48 [0.29-0.77], p=0.003), versus being alone (reference category) (**Figure 5B and Table 2**).

**Table 2.**
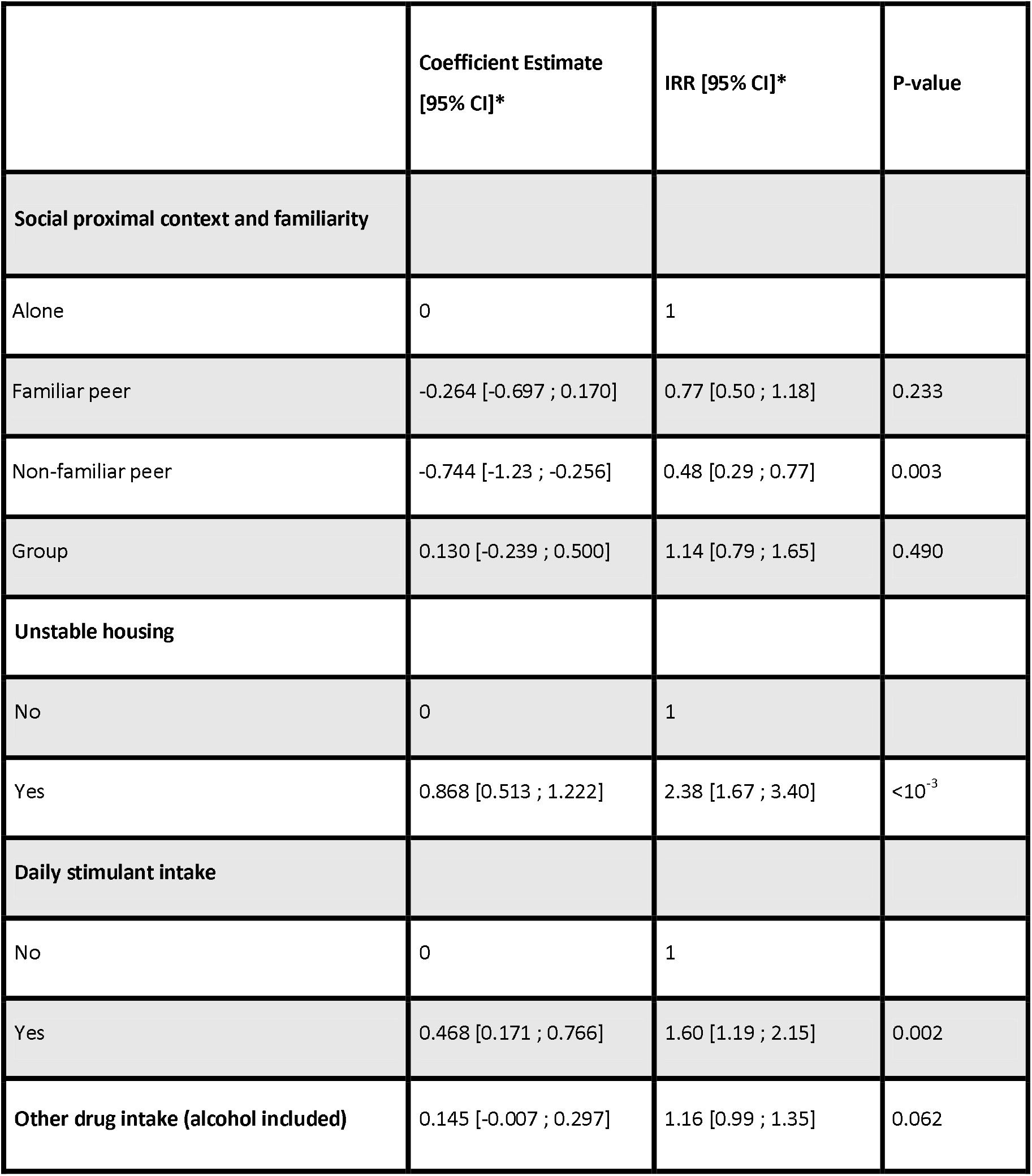
**Human study** - Association between familiarity / peer presence and cocaine use frequency among humans (N=77, 246 episodes of cocaine use) – multivariable analysis.

|  | Coefficient Estimate<br>[95% CI]* | IRR [95% CI]* | P-value |
| --- | --- | --- | --- |
| **Social proximal context and familiarity** |  |  |  |
| Alone | 0 | 1 |  |
| Familiar peer | -0.264 [-0.697 ; 0.170] | 0.77 [0.50 ; 1.18] | 0.233 |
| Non-familiar peer | -0.744 [-1.23 ; -0.256] | 0.48 [0.29 ; 0.77] | 0.003 |
| Group | 0.130 [-0.239 ; 0.500] | 1.14 [0.79 ; 1.65] | 0.490 |
| **Unstable housing** |  |  |  |
| No | 0 | 1 |  |
| Yes | 0.868 [0.513 ; 1.222] | 2.38 [1.67 ; 3.40] | <10<sup>-3</sup> |
| **Daily stimulant intake** |  |  |  |
| No | 0 | 1 |  |
| Yes | 0.468 [0.171 ; 0.766] | 1.60 [1.19 ; 2.15] | 0.002 |
| **Other drug intake (alcohol included)** | 0.145 [-0.007 ; 0.297] | 1.16 [0.99 ; 1.35] | 0.062 |

Other correlates associated with greater frequency of cocaine use in the multivariable analysis were unstable housing (IRR [95% CI]=2.38 [1.67-3.40], p<10^−3^), daily stimulant use (IRR [95% CI]=1.60 [1.19-2.15], p=0.002) and other substances (alcohol included) concomitantly used during the episode (IRR [95% CI]=1.16 [0.99-1.35], p=0.062).

## Discussion

Our study showed that the presence of a peer at the time of stimulant intake has a beneficial reducing effect on stimulant consumption. Furthermore, these effects are modulated by both the familiarity and the former drug-using status of this present peer. Indeed, the lowest level of consumption was observed in presence of a non-familiar peer naïve of cocaine. All these results were replicated when rats were in the presence of a peer also allowed to take cocaine.

Validating our model, these beneficial effects of the peer presence were also observed in humans, as well as the modulating effect of the peer’s familiarity status. **We also addressed in this paper the issue of neurobiological basis of social influence on drug consumption and showed a particular role for STN**. Indeed, reducing STN activity could be even more efficient to potentiate the beneficial effect of the presence of a peer during drug consumption. In fact, in contrast with the sham-control, STN-lesioned rats decreased their consumption in presence of a peer, whatever the familiarity and the former drug using status of the peer, revealing that STN is involved in social processes, as confirmed in the social preference experiments.

**Finally, we developed and tested in this article a novel approach in terms of design and statistical analysis to conduct translational research** on the influence of proximal social factors on a standardized outcome. This could have important repercussions in research on human behaviors and may encourage other behavioral researchers to adopt a similar approach especially when the research question can be translated into public health actions.

### General influence of social presence

The first result of this study is that, in line with former results^28^, the presence of a peer during cocaine self-administration has beneficial effects on stimulant consumption, supporting the hypothesis that the rewarding properties of social contact may outbalance the reinforcing properties of drug consumption^25,26^ and modulate the affective valence of drug use^52^. This explanation is concordant with the results of the conditioned place preference test we performed where rats (housed in pairs) preferred an environment where they were in contact with their home-cage partner over an environment where they remained alone. Interestingly, and more closely to the common drug use situations reported in human subjects, when the present peer could also take drug, we also observed a decreased consumption. This is opposite to what has been reported in rats acquiring the drug self-administration simultaneously^53^. In the present study, the animals had acquired the behavior separately and were only tested simultaneously when they had reached a stable intake. This major difference may possibly account for the opposite influence observed here and suggests that social context can be detrimental during acquisition of a behavior but can help reduce it when well established.

### Influence of peer familiarity and dominance status

We have shown that sham-control rats decreased more their consumption in presence of a stranger peer than in presence of a familiar one, when this peer was an observer. Following the hypothesis above, this result suggests that the presence of a stranger would be more rewarding than that of a familiar peer. In line with this, results of the social preference tests showed that sham-control rats exhibited a preferential exploration for the social stimulus over an object, as expected^54^, but more interestingly, the effect was stronger when the social stimulus was a stranger rat than when it was a familiar peer. Furthermore, in the CPP experiment, the presence of the cage-mate appeared to be rewarding only for the dominant animal, but not for the subordinate. As a result, it was thus coherent to observe that the presence of the cage-mate only decreased the cocaine consumption for the dominant rats, but not for the subordinates being observed by their dominant peer. Altogether, our results show that, on one hand, the presence of a stranger peer is highly rewarding for rats and leads to decreased cocaine intake. On the other hand, the presence of a familiar peer (i.e: the cage-mate) is also rewarding and leads to decreased stimulant consumption, but only for the dominant rat.

An alternative explanation for the difference between familiar and non-familiar peer and their influence on self-administration behavior could be that the presence of the latter represents a powerful distractor^55^. A subject’s attention may be focused on the non-familiar peer rather than on the drug, consequently leading to decreased frequency of drug consumption. A recent study in monkeys has shown that the presence of a peer increases the activity of attention-related cerebral structures^56^. Moreover, in line with this hypothesis, it has also been shown by Huguet and colleagues^57^ in baboons that social presence, especially if potentially threatening (dominant or non-familiar) could divert attention from the focal task and also consume cognitive control resources. The level of stress induced by the dominance status might play a critical role in this effect as well. It would be interesting to measure the level of cortisol in our animals; although the monkey study has shown that the presence of peer did not modulate cortisol levels^56^.

Interestingly, when the peer could also self-administer cocaine, its familiarity status did no longer influence the consumption of the subject. Those results suggest that the effects of peer drug-using status may outweigh those of its familiarity status, highlighting the importance of considering the different characteristics of proximal social factors to better understand their mechanisms.

### Influence of peer drug-using status

Sham-control rats decreased their cocaine consumption more in presence of a peer which was not immediately consuming than when in presence of a consuming peer. Furthermore, this decrease was higher when the peer was naïve to the cocaine, highlighting the modulatory effect of the drug-using status of a peer present during cocaine self-administration^52^. In line with our results, an econometric analysis has shown that the reinforcing properties of cocaine diminished when the peer present was abstaining^58^. This effect could rely on social-learning theories of substance-use, suggesting that people in a group of drug-users tend to imitate each other^58^. Illustrating this theory, Smith et al. have shown that the presence of a rat also self-administering cocaine increases drug consumption during simultaneous acquisition^53^, as discussed above.

### The same influence of social proximal factors in rats and humans?

Although former epidemiological studies have shown the importance of relationships between peers immediately during drug consumption and on the sharing of injecting equipment, drug seeking^59^ and craving^59–62^, our experiment is the first studying the nature of the relationship within a dyad of drug-users and to correlate it with cocaine consumption. Results on humans mirrored the effect of the familiarity status found in our rat model, since like in rats, the presence of a peer decreased cocaine intake, and this decrease was higher if the peer was stranger. On the other hand, we were not able to estimate the effect of the peer drug-using status in humans, since in all drug consumption episodes reported here, the present peer were also a consumer.

Regarding rat and human results, we have to note some limitations in our study. First, while rats study design was experimental, the human study collected retrospective information on episodes of stimulant consumption. In the latter, outcomes may be subject to recall and social desirability biases and too confounding. Nevertheless, we questioned participants about their most recent episodes of drug intake to minimize recall biases. Furthermore, as cocaine use may vary across the different episodes, measuring use only from the most recent episode allowed us to simultaneously record the frequency of cocaine consumption and the relationship with a present peer.

Despite these limitations, we found very similar estimates for the association between consumption frequency and the presence of a peer, with a modulatory role of the familiarity of this peer in both rat and human models.

### The STN as a substrate of the influence of social proximal factors on drug intake

Our results show that STN modulates the influence of the presence of a peer on cocaine intake. STN is a part of the basal ganglia and is known to be involved in motivation and addiction-related behaviors^36,63–65^. Interestingly, our results show that the baseline consumption of STN-lesioned and sham-control rats is similar. This is in line with former studies showing that STN lesion modulates cocaine intake in certain conditions (increased motivation in progressive ratio^36,47,66^, escalation under FR1 or FR1 after abstinence^64^). On the contrary, in STN-lesioned rats, the presence of a peer induced a strong decrease in cocaine intake (equivalent to that induced by a stranger peer presence in sham control rats). Interestingly, this decrease was not modulated by familiarity, the dominance status, nor the drug-using status of the peer.

Furthermore, results of social preference tests showed that STN-lesioned rats had the same preferential exploration for a stranger or the cage-mate rat, over an object. Likewise, both dominant and subordinate STN-lesioned rats showed a preference for their cage-mate in the CPP experiment. Altogether, these results suggest that in STN-lesioned rats, familiarity and dominance status no longer modulate the rewarding value of a peer presence. As a result, in STN-lesioned rats the presence of a peer decreases cocaine intake, independently of its familiarity or dominance status. This suggests that they could no longer make the difference between familiar and non-familiar peers. Interestingly, in humans, in line with our results, STN-DBS in parkinsonian patients has been shown to blunt facial and vocal emotion recognition^37,67–69^. Deficits in emotional processing have also been observed in STN-lesioned rats^38^, notably in emotional communication^39^. Then, the involvement of the STN in social-emotional processes could explain the blunted differential rewarding value of the presence of a peer depending of their characteristics observed in STN-lesioned rats. However, the involvement of the STN in social cognition remains to be further investigated.

To conclude, this study provides three major contributions. First, we have shown that peer presence, familiarity and drug-using status of this peer have major effects on drug consumption. Those characteristics must be taken into account to ameliorate existing models in the study of the influence of proximal social factors on drugs consumption. Second, we have shown parallel influences of proximal social factors on cocaine use in rats and humans, notably regarding the effect of the familiarity status of the peer present. This parallel highlights translational potential from rats to humans. The need for translational studies^70,71^ is essential for a better understanding of the proximal social factors influence on addiction^72^, notably their neurobiological substrates. Finally, we have shown that STN appears to play a critical role in the influence of those factors. Modulating its activity may thus result in an alteration of social context influence on drug use, toward decreased cocaine consumption. Since STN deep brain stimulation is proposed as a therapeutic strategy to treat addiction^66^, this apparent emotional side-effect could serve to reduce further the drug intake, as seen here in STN lesioned rats.

Understanding how proximal social factors modulate drug consumption will help in the design of novel preventive and therapeutic strategies including social interventions to target drug-using populations. Furthermore, the presence of a non-familiar and possibly drug-naive peer would appear to be a driver for diminished stimulant intake. Given that there is still space for improvement in the management of cocaine-related disorders, these results may be crucial to develop harm reduction strategies for stimulant users. At the clinical level, this would translate into involving peers in treatment education. At the health policy level, it would mean promoting the use of harm reduction strategies, such as peer education on injection and the deployment of supervised consumption rooms.

## Acknowledgements

Our thanks to Dany Palleressompoulle and Joel Baurberg for their technical support, as well as to the animal facilities’ personnel. Our thanks also to Silvia Rosellini and Chiara Calzolaio for help with statistical analyses and to Jude Sweeney and Maya Williams for the English revisions and editing of the manuscript. This research was funded by CNRS, Aix-Marseille Université (AMU), the “Agence Nationale pour la Recherche” (ANR_2010-NEUR-005-01 in the framework of the ERA-Net NEURON to C.B. and supporting Y.P.), the Fondation pour la Recherche Médicale (FRM DPA20140629789 to C.B.), and the support of the A*MIDEX project (ANR-11-IDEX-0001-02) funded by the “Investissements d’Avenir” French Government program, managed by the French National Research Agency (ANR).

## Authors contribution

E.G, C.V conducted most of the rats experiments, with the help of L.G, C.M

S.N, PR conducted the human experiments

C.B, C.V, E.G, P.C and S.N, wrote the paper.

V, C.P, C.V, E.G, S.N, P.R, conducted the statistical analysis.

C.M, C.V and E.G conducted the histological analysis.

C.B, C.V, E.G, K.D, P.H and Y.P designed the rat experiment

CB, PR, PC and SN designed the human questionnaire

CB, PC and PH obtained the funding from A*MIDEX

## Materials and Methods

### Rat Study

#### Animals & surgery

90 male Lister Hooded rats (Charles River Laboratories, Saint-Germain-sur-l’Arbresle, France) were used in the present study. They were subjected to STN lesion procedure and then 64 of them were subjected to an intravenous silicon catheter implantation for the self-administration study (see Supplemental Materials for more detailed procedure).

#### Apparatus

All the drug self-administration experiments were conducted in four custom-built self-administration chambers (60×30×35cm) divided into two compartments, separated by a grid. (See Supplemental Materials for more details). A picture and a schematic diagram of the self-administration chambers are shown in **Figure 1**.

As for the social experiments, they were conducted in a rectangular plastic arena (70×30×40cm, social preference test) and opaque Perspex boxes (90×35×33cm, conditioned place preference). (See Supplemental Materials for more details).

#### Experimental Procedure

##### Influence of the presence of peer and its characteristics on cocaine self-administration in sham and STN lesioned rats

2 sets of animals were used in this first experiment. One set of animals (sham n=14, STN lesioned n=17, experiment “observing peer”) were subjected to the self-administration procedure while a peer was present, just observing, while other animals (sham, n=14, experiment “co-self-administering peer”) were allowed to take cocaine in presence of another peer also self-administering cocaine.

Both sham and STN lesioned rats were individually trained to pull a chain to self-administer cocaine (80 µg per 90 µl infusion in 5s, 0.2 mg/kg) under a continuous schedule of reinforcement (Fixed Ratio 1 (FR1), 1 chain pulling resulted in 1 cocaine injection) for daily one-hour sessions. Since chain pulling is not as easy as pressing a lever, baseline level of cocaine consumption was too low with the classic dose of 250 µg per 90µl. We, thus, chose to decrease the dose of cocaine to 80 µg to increase the number of injections taken per session. For half of each group, cocaine was randomly assigned to one of the two chains (the active chain) and the other half of the group had the opposite rule. Pulling the active chain switched on the cue-light, delivered the cocaine to the blood stream and started a 20-s “time-out” during which any further pulling was recorded as perseveration, but had no other consequence. Pulling the other chain (inactive chain) was also recorded, as an error, and had no consequence. Once consumption became stable (i.e. 5 consecutive days with the same number of cocaine injections ±2), the last 5 days of acquisition were used as a baseline for cocaine consumption when the rats were alone in the apparatus (condition “alone”). For each one-hour behavioral session, the number of cocaine injections was recorded.

Cocaine self-administration with an observing peer

The rats were exposed to 4 different conditions, for 5 consecutive days each, each session lasted 1 hour:

1. Alone (baseline consumption): The rats were alone in the self-administration chamber during acquisition of the self-administration behavior. Once they have learned it, the last 5 sessions were kept for analysis.
2. Peer presence: The self-administering rats were in presence of another rat (hereafter “peer”), having no access to cocaine, present as an observer. This peer could be for some rats “familiar” (i.e. a cage-mate also trained for self-administration; (sham n=8 STN lesioned n=6) or “stranger” (hereafter “non-familiar peer”) (i.e. a rat trained for cocaine self-administration but living in a different home-cage; sham n=6 STN lesioned n=11). Observing peers were introduced into the cage after they had a minimum of 4 hours of abstinence from cocaine. For the “familiar peer” condition, the dominant status (“dominant” observed by “subordinate” vs “subordinate” observed by “dominant” peer) was also taken into account, as it could be determined after behavioral assessment described in Supplemental Materials.
3. Post-peer presence: Rats were “back alone” after exposure to peers (sham n=14, STN lesioned n=17) in order to assess whether peer presence could have a long-lasting influence on cocaine intake even in absence of this peer.
4. Non-familiar and cocaine-naive peer presence: rats were tested in presence of a rat from another group that had never been exposed to cocaine (sham n=11 STN lesioned n=15).

The same peer was used for each given rat for the 5 behavioral sessions of a condition.

Cocaine self-administration with a co-self-administering peer

During this part of the experiment, in the peer presence condition, both rats could have access to cocaine during the 1-hour behavioral sessions. Acquisition of the self-administration behavior was done in “alone” condition. Then, rats were subjected to 2 conditions:

1. Alone (baseline consumption): when the rats were alone in the self-administration chamber.
2. Peer presence: In the presence of another rat (hereafter “peer”) also having access to the drug, so the rats could both have access to cocaine during the one-hour session. This peer could be either “familiar” (i.e. a cage-mate also trained for self-administration; n=6) or stranger (hereafter “non-familiar peer”) (i.e. a rat trained for cocaine self-administration but living in a different home-cage; n= 8).

The same familiar and non-familiar peers were used for each rat for all behavioral sessions.

#### Reinforcing properties of Social presence

##### Social preference

To examine if the STN is involved in social preference and could modulate rat’s social preference for a social stimulus (a peer) versus an object, we subjected rats to a social preference test ^44,45^. In this procedure, the subject rat is simultaneously exposed to 2 identical cages, one containing an object and the other a social stimulus (either a stranger rat or the cage-mate). Subject rat is allowed to freely explore for 5 minutes both stimuli. Investigation time has been defined, as previously described in Engelmann (1995)^46^, as the time spent by the subject actively exploring (sniffing with the tip of the nose within approximately 1 cm of the cage containing the stimulus rat or the object. To reduce the number of animals, the same cohort was used for both familiar and stranger stimulus (sham-control rats n=7 and STN-lesioned rats n=9). The order of testing “familiar” or “stranger” stimuli was counterbalanced to prevent a possible effect of repetition of the testing. The objects were changed for each session. The investigatory behavior has been scored by a trained observer. For every test, if the total investigation time was less than 30-sec, the results were removed from the analysis.

##### Conditioned Place preference

In order to assess whether or not cage-mate presence can be rewarding for a rat, conditioned placed preference to an environment associated with the presence of the cage-mate was performed. As previously described^47^, the CPP procedure lasted 10 days. Each session was video recorded for further analysis. On the pre-conditioning day (Day 1), 66 sham rats and 14 STN lesioned rats were placed in the CPP apparatus and allowed to explore the two compartments for 15 minutes. The time spent in each compartment was recorded manually in seconds to determine a possible natural preference for each rat. During conditioning, rats were exposed for 30 min either to their cage-mate in their less-preferred compartment on days 2, 4, 6 and 8 or to no other rat in their initially preferred compartment on days 3, 5, 7 and 9. On day 10, all rats were placed in the middle of the CPP apparatus for 15 minutes and both compartments were made accessible for exploration. The time spent in each compartment was recorded manually in seconds and the score of preference for the compartment associated with the presence of the cage-mate was calculated.

#### Statistical analysis

To analyze the self-administration experiments, we assessed the number of cocaine injections during the one-hour cocaine self-administration sessions (i.e. frequency; outcome) with regard to several parameters (see Supplemental Materials for more details).

For the social preference test, all variables are expressed as mean number ± SEM and the p-value threshold have been set at α=0.05 (See Supplemental Materials for more details).

Concerning the conditioned place preference results, we analyzed the preference score of the rats for the compartment associated with the stimulus (see Supplemental Materials for more details).

### Human study

#### Design

The human study is a cross-sectional survey (DDYADS) implemented between October 2015 and June 2016 in five cities in France characterized by high prevalence of drug use (Marseille, Paris, Montreuil, Saint Denis, Nice).

#### Participants

Seventy-seven French-speaking regular stimulant users – defined as using cocaine or methylphenidate ≥5 times a month – were recruited in different sites between October 2015 and june 2016. The study received authorization from the national French Data Protection Authority (CNIL) and Aix-Marseille University’s institutional review board. All participants provided written informed consent.

#### Data collection

Data were anonymously collected through a face-to-face standardized questionnaire administered by trained interviewers.

Social environment at the moment of stimulant use was described as follows: alone, with one peer (“familiar” or “non-familiar”), with a group (i.e., two or more peers). For episodes involving the participant and one peer, information on the peer was collected (see Supplemental Materials for more details).

#### Outcome

The outcome was the number of times drugs were consumed in one hour during each episode (i.e. frequency of drug intake).

The frequency of use during one episode (standardized by the duration of the episode) is an interesting measure, especially among cocaine users where the half-life of the drug may require repeated intake. From a public health viewpoint, this frequency is associated with several health risks (e.g. overdoses and other fatal events, as well as unsafe sexual behaviors^48–50^).

#### Statistical analysis

We considered each drug intake episode reported during the interview as a statistical unit. In order to take into account the within-subject correlation due to repeated measures (i.e., drug use episodes in the previous month) reported by the same individual, we used the Poisson Generalized Estimated Equations (GEE) approach for count data^51^ (see Supplemental Materials for more details). Both Poisson mixed models and GEE provide, for each predictive/explanatory variable associated with the frequency of cocaine/methylphenidate intake, an estimate of the incidence rate ratio (IRR) or relative risk and its 95% confidence interval (CI). Confidence intervals not containing 1 indicate a significant association (p-value<0.05). IRRs are a measure of the association between the explanatory variable and the frequency of cocaine intake.

## Supplemental material

### Translational issues

The challenges in combining the results of the two studies were overcome thanks to several multidisciplinary sessions which included epidemiologists, neurobiologists/psychologists and statisticians. Working together, these stakeholders ensured the validity of both studies’ designs, and decided “a priori” on the strategic statistical analysis plan to implement in order to facilitate comparison.

More specifically, the three challenges overcome concerned: 1) design 2) explanatory variables 3) outcomes and statistical analysis.

With respect to the first challenge concerning design, it should be noted that cocaine administration can be randomized in rats but not in humans, at least in France. We therefore used a randomized plan (see experimental design section for rats) for the former, but for ethical concerns, an observational cross-sectional epidemiologic design for the latter. This observational design allowed us to retrospectively explore each episode of cocaine/methylphenidate consumption and the associated social context. We adapted a methodology already used in network analysis by Buchanan and Latkin^6^. It is important to underline that the experimental conditions for humans (i.e., intake in the presence or not of peers, intake with or without familiar, subordinate or cocaine-naïve peers) mirrored the experimental conditions in the study for rats.

The second major challenge was to “translate” the concepts of familiarity, subordination among peers and cocaine naivety from rats to humans. Due to the lack of literature on this specific issue, this “translation” was constructed using open-ended questions which explored these same three concepts. This approach helped to provide an operational definition of what familiarity, subordination, and cocaine naivety mean for cocaine-using peers.

For the third challenge, we used the same outcome (“frequency of cocaine injection/duration of the session/episode”) and the same models in both studies to provide both an adequate estimate of the association between each “social context” and the outcome. Poisson regression models (based on generalized estimating equations for humans and mixed models for rats) were used to obtain comparable estimates of the associations found between each social context and the frequency of cocaine intake in rats vs. humans.

## Supplemental Material and methods

### Rat study

#### Animals & Surgery

90 male Lister Hooded rats (Charles River Laboratories, Saint-Germain-sur-l’Arbresle, France) were housed in pairs upon their arrival. Rats were handled 2-3 times a week. They were maintained under 12-h light/dark cycles and had ad libitum access to food and water. All animal care and use conformed to the French regulation (Decree 2010-118) and were approved by local ethic committee and the ministry of agriculture (#3129.01). Rats (n=90) were subjected to the STN lesion or sham intracerebral injection procedure. Then, using standard surgical procedures, silicon catheters were inserted into the right jugular vein of the rats (n=64). Due to mislocation of the lesion and the difficulty to maintain catheter, 19 rats in total were excluded from the study. Among them, 10 were discarded from Experiment 1 (i.e. the double self-administration experiment. When one rat could no longer be tested (catheter patency issue), its peer had to be excluded as well). The histological illustration of the lesion is shown in **Figure S1** (see supplemental Material).

Rats were anesthetized with ketamine (100 mg/Kg, s.c., Imalgène 1000, Merial, Lyon, France) and medetomidine (0,2 mg/Kg, s.c., Domitor, Orion Pharma, Espoo, Finland). At the end of surgery, medetomidine was reversed by 0.2 mg (4.28 mg/Kg, s.c.) atipamezole (Antisedan, Orion Pharma, Espoo, Finland).

##### Catheter implantation

Using standard surgical procedures, silicon catheters were inserted into the right jugular vein of the rats and exited dorsally between the scapulae.

From this point forward, catheters were daily flushed with 0.2 ml of a sterile saline solution containing heparin (Heparin Sodium, Sanofi, Paris, France; 5000 U.I/ml) and enrofloxacin (Baytril 5%, Bayer, Loos, France; 50 mg/ml). Catheters were also regularly tested with propofol (Propovet, Abbott, 10 mg/ml) to confirm their patency.

##### Subthalamic nucleus lesion

Rats were secured in Kopf stereotaxic apparatus. Then, unilateral 30-gauge stainless-steel injector needles connected via Tygon tubing (Saint Gobain performance plastics) to a 10µL Hamilton microsyringe (Bonaduz, Switzerland) fixed on a micropump (CMA, Kista, Sweden) were positioned into the STN. Coordinates for the aimed site were (with tooth bar set at -3.3mm): anteposterior -3.72 mm; lateral ±2.4 mm from bregma; dorsoventral -8.4 mm from skull (Paxinos & Watson, 2005). Rats received either a bilateral injection of ibotenic acid (9.4µg/µL, AbCam Biochemical, Cambridge, UK; STN-lesioned group n=50) or vehicle solution (phosphate buffer, 0.1M; Sham control group n=40). The volume injected was 0.5µL per side infused over 3 min. The injectors were left in place for 3 min to allow diffusion. At the end of surgery, medetomidine was reversed by 0.2 mg (4.28 mg/Kg, s.c.) of atipamezole (Antisedan, Orion Pharma, Espoo, Finland).

#### Procedures

##### Influence of the presence of peer and its characteristics on cocaine self-administration in rats

###### Apparatus

All the self-administration experiments were conducted in four custom-built self-administration (SA) chambers (60 cm x 30 cm x 35 cm) made of opaque Perspex and divided into two compartments, separated by a grid (2.5 cm x 2.5 cm). For the first part of the experiment (simple self-administration), only one of the two compartments was equipped with 2 chains and a stimulus light located on the right-hand wall. For the second part of the experiment (double self-administration), both compartments were equipped so that both rats could self-administer. The grid allowed each rat visual, auditory and olfactory communication, limited tactile contact with its peer, and prevented each rat from accessing the tethering system of its peer. Drug infusions were delivered via intrajugular route of administration through tubing protected by a stainless steel spring, connected to 10 ml cocaine syringes positioned on motorized pumps (Razel Scientific Instruments, St. Albans, VT, USA) outside of the chamber.

All the chambers and pumps described above were controlled by a custom-built interface and associated software (built and written by Y. Pelloux).

###### Data analysis and statistical analysis

We assessed the number of cocaine injections during the one-hour cocaine self-administration session (outcome; i.e. frequency) with regard to the state of the STN (intact i.e. sham rats or lesioned), the nature of the social relationship with rat peers (familiar or not familiar), the history of cocaine exposure of rat peers (naive or non-naive) and whether the rat peer was dominant or subordinate.

Poisson mixed models were used to take into account the correlation over time between repeated measures of the outcome.

The following four models were analyzed for sham rats and then for STN lesioned rats, each including a random effect on time (in days), and the following experimental factor:

A. Peer presence: alone or with a non-naive peer, irrespective of the relationship (i.e. familiar/not familiar)
B. Familiarity: alone, with familiar peer, with non-familiar peer (only non-naive peers)
C. History of cocaine exposure: alone, with naive peer, with non-naive peer, back alone
D. Social status: dominant or subordinate (only non-naive familiar peers)

Then the following analysis were done to take into account the difference between sham and STN lesioned rats, each including a random effect on time (days), and the following experimental factor:

A. Baseline consumption: sham or STN lesioned when alone
B. Peer presence: Sham or STN lesioned (non-naïve and naive peers)

##### Reinforcing properties of Social presence

###### Apparatus

The social preference test took place in a rectangular plastic arena (70 cm length, 40 cm high, 30 cm depth), with a fresh litter, changed between each test. Stimulus rats has been habituated for 10-min in the cage before the test, to avoid a fear reaction. Every exposure of the subject rat with a stimulus has been recorded with a webcam connected to a computer, using Bonsai (Open Ephys).

Concerning the conditioned place preference experiment was performed in two opaque Perspex boxes (90 cm length x 35 cm width x 33 cm height) divided into two main compartments - A and B (45 × 35 × 33 cm each). Compartments A and B differed in the spatial organization of 2 columns. During conditioning, the compartments were separated by an opaque screen to confine the animals to one or the other. During habituation and testing, the screen was removed, allowing the animals to explore the entire apparatus. All the social preference tests were run in a quiet and dimly illuminated experimental room. Arenas, objects and cages were cleaned between each test, using a solution of hydrogen peroxide.

###### Data analysis and statistical analysis

For the social preference test, all variables are expressed as mean number ± SEM and the p-value threshold have been set at α=0.05

Analyses and graphs were carried out using Prism (GraphPad). Time spent exploring the different stimuli was compared between sham-control and STN-lesioned rats using Wilcoxon matched-pairs signed rank test.

Concerning the conditioned place preference results, we analyzed the preference score of the rats for the compartment associated with the stimulus. This score was calculated as the time spent in the environment previously paired with the social interaction (test day), minus the time spent in the same compartment before it was associated with the reward (pre-conditioning day). This gave us an indication of the positive or negative memory that the animal had of its social experience. The non-parametric Mann-Whitney test was used for the conditioned place preference experiment to assess the group effect while the Wilcoxon signed rank test was used for the conditioning effect.

##### Dominance status assessment

For the self-administration and CPP studies, dominance status within each pair of rats was assessed by behavioral observation during the first conditioning session of the CPP experiment. The number of pinning and pouncing for each rat was recorded during a 15-min period of interaction. The “dominant” rat was assumed to be the one doing the most pinning and pouncing. The other rat was qualified as “subordinate”.

#### Histology

The cresyl violet staining allows visualization of the consequences of the ibotenic acid injection in the STN that are characterized by glial reactivity and neuronal loss. Thus, after verification, rats presenting a non-sufficient lesion or a misplaced lesion were discarded from the analyses (**Figure S1**).

### Human study

#### Participants

Seventy-seven French-speaking regular stimulant users – defined as using cocaine or methylphenidate ≥5 times a month – were recruited in different sites, including methadone centers, harm reduction centers, low-threshold mobile health units, and through word-of-mouth referrals, between October 2015 and June 2016. Non-prescribed methylphenidate was also considered as cocaine users may switch from cocaine to methylphenidate and vice-versa in the areas where the study was conducted, depending on black market availability and costs. Table S1 describes participant characteristics (N=77) and details of drug use episodes involving intranasal and intravenous routes of administration (246 episodes).

#### Data collection

Data were anonymously collected through a face-to-face standardized questionnaire administered by trained interviewers. Interviews were conducted in a dedicated room at centers or in a café and lasted from 30 to 60 minutes. Participants were remunerated with a €15 gift voucher for completing the interview.

To minimize recall bias, participants retrospectively described episodes during the previous month where they used stimulants.

Social environment at the moment of stimulant use was described as follows: alone, with one peer, with a group (i.e., 2 or more peers). For episodes involving the participant and one peer, information on the peer was collected.

Peers were considered “familiar” if they were close friends or relatives, and if the participant could speak about his/her intimate life with them. Otherwise, peers were considered “non-familiar”. Participants were considered subordinate if they were economically dependent on the peer or if the peer was the leader in terms of drug use contexts (e.g. paying for the drug). Each drug use episode was characterized as follows: principal route of administration (intravenous, intranasal), type of stimulant (cocaine or methylphenidate), drug effect perception (from 1 to 5), concomitant use of other psychoactive substances including alcohol, location where episode took place (public versus private), state of mind (positive versus neutral versus negative), number of times drugs were consumed (including alcohol), and episode duration. We also collected data on participant characteristics including age, gender, employment status, educational level, housing situation (stable versus unstable), hazardous alcohol use (AUDIT-C score) (Bush & al., 1998), financial problems, including economic dependence on the peer, and the number of days the participant had used stimulants in the previous month.

#### Statistical analysis

We considered each drug intake episode reported during the interview as a statistical unit. In order to take into account the within-subject correlation due to repeated measures (i.e., drug use episodes in the previous month) reported by the same individual, we used the Poisson Generalized Estimated Equation (GEE) approach for count data^51^.

First, we conducted a univariable analysis to test each variable describing the “social context”-peer presence (alone vs. one peer vs. group), familiarity, dominance of the peer – and potential correlates/confounders.

Second, based on the results of the univariable analysis, two models were built. In the first, we tested the role of peer presence (alone, with one peer, with a group i.e., two or more peers) on the frequency of stimulant use, after taking into account potential correlates/confounders including: 1) context of the stimulant use episode: type of location (private versus public place), route of administration (intravenous, intranasal), state of mind (positive, neutral, negative), number of other substances concomitantly used (including alcohol); 2) participant characteristics: gender, age, educational level (< high school certificate versus ≥ high school certificate), employment status, stable housing (i.e. renter or owner of their personal housing versus other), financial difficulties, hazardous alcohol use (AUDIT-C ≥4 for men and ≥3 for women), and number of days of stimulant use during the previous month (daily stimulant user versus other).

The second model was built to examine the role of peer familiarity (alone, with one familiar peer, with one non-familiar peer, with a group) on the frequency of drug intake after taking into account the potential correlates/confounders described in the first model.

These models provide, for each predictive/explanatory variable associated with the frequency of cocaine/methylphenidate intake, an estimate of the incidence rate ratio (IRR) or relative risk and its 95% confidence intervals (CI). Confidence intervals not containing 1 indicate a significant association. IRRs are a measure of the association between the explanatory variable and the frequency of cocaine intake. For example, a significant IRR of 0.30 for the presence of a peer compared with being alone means that the relative risk of intake of cocaine over the duration of the episode decreases by 70% when an individual (rat or human) is with a peer. The use of these IRRs allows comparison of the association found between each social context and the frequency of cocaine use in both rats and humans.

We used a liberal p-value<0.20 in the univariable analysis to identify social context explanatory variables eligible for each multivariable model. A backward selection procedure was used to determine the two final multivariable models. We set the p-value threshold at α = 0.05 for these latter. STATA/SE version 12.1 software for Windows was used for the analyses.

#### Data availability statement

The datasets generated during and analyzed during the current study are available from the corresponding author on reasonable request.

##### Description of Human Study Group’s Characteristics

Table S1 describes participant characteristics (N=77) and details of drug use episodes involving intranasal and intravenous routes of administration (246 episodes).

More than 80% of the study samples were males with median age of 41 years. By definition they were all cocaine users and more than one third reported daily use. Median age at first use was 17 years. Most reported polysubstance use. Two thirds of the study sample reported either exclusive cocaine use or with methylphenidate. Approximately half were classified as hazardous alcohol users and 45% as heavy smokers (more than 20 cigarettes/day). One third reported having a high school certificate and 42% declared stable housing. The majority (66.2%) reported financial problems.

Concerning the median [interquartile range (IQR)] frequency of stimulant consumption was 1.05 [0.05-2.6] when subjects were alone and 1 [0.3-2] when with peers. This study group reported 246 episodes (on average 3.2 episodes per person) of stimulant use alone (26.8%), with a peer (48.8%) or in a group (24.4%). The majority of episodes reported by the study group occurred in presence of a familiar peer (29.4%), followed by alone (26.9%) and within a group (24.5%) while episodes of stimulant use alone accounted for 19.2%. It is worth noting that in 67.1% of the reported episodes, study participants had injected the drug, and that most episodes of stimulant use occurred in a public space. The majority of episodes occurred in the afternoon and the median number of substances used (including alcohol) at any episode was 2.

---

**Table S1.**
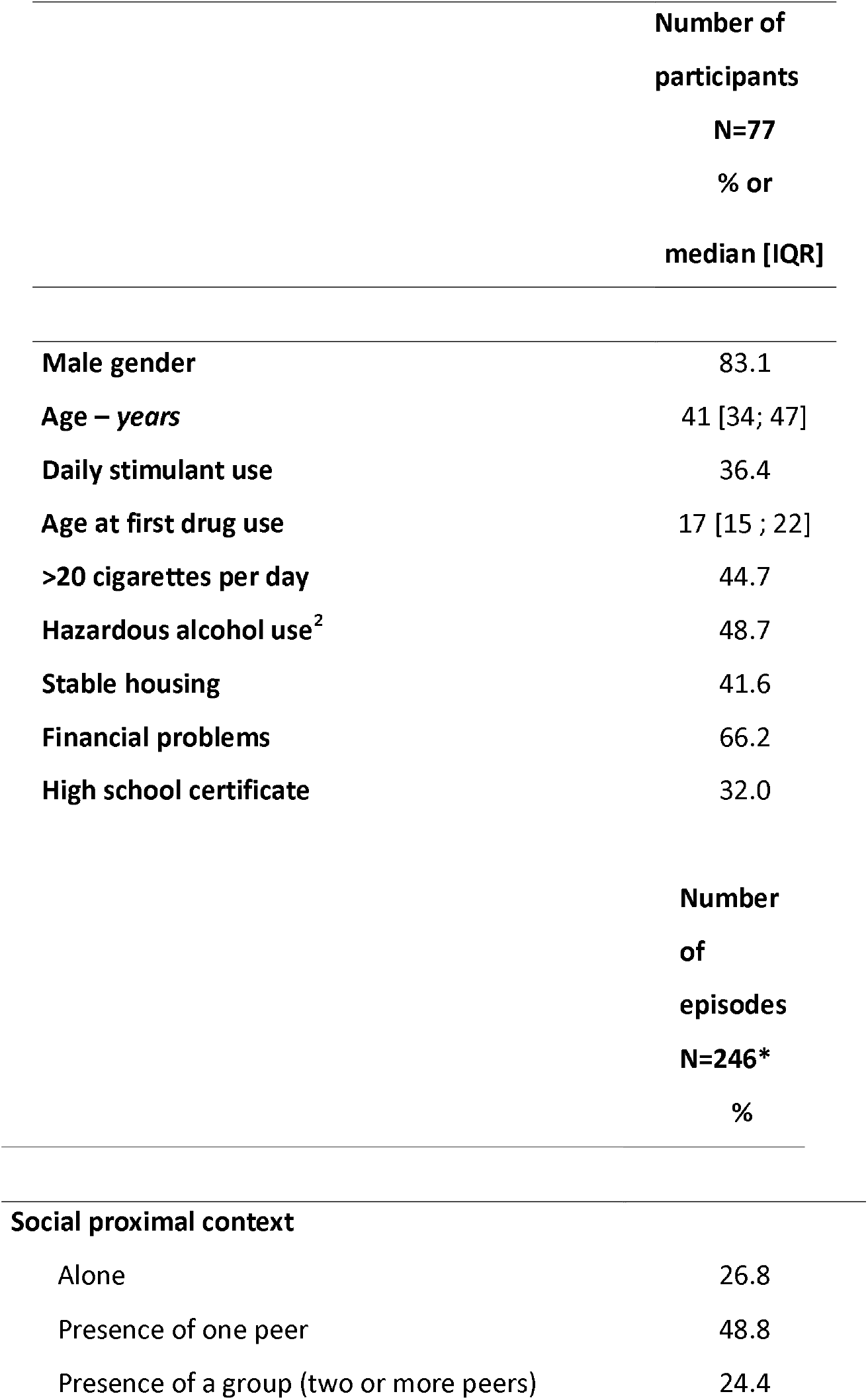
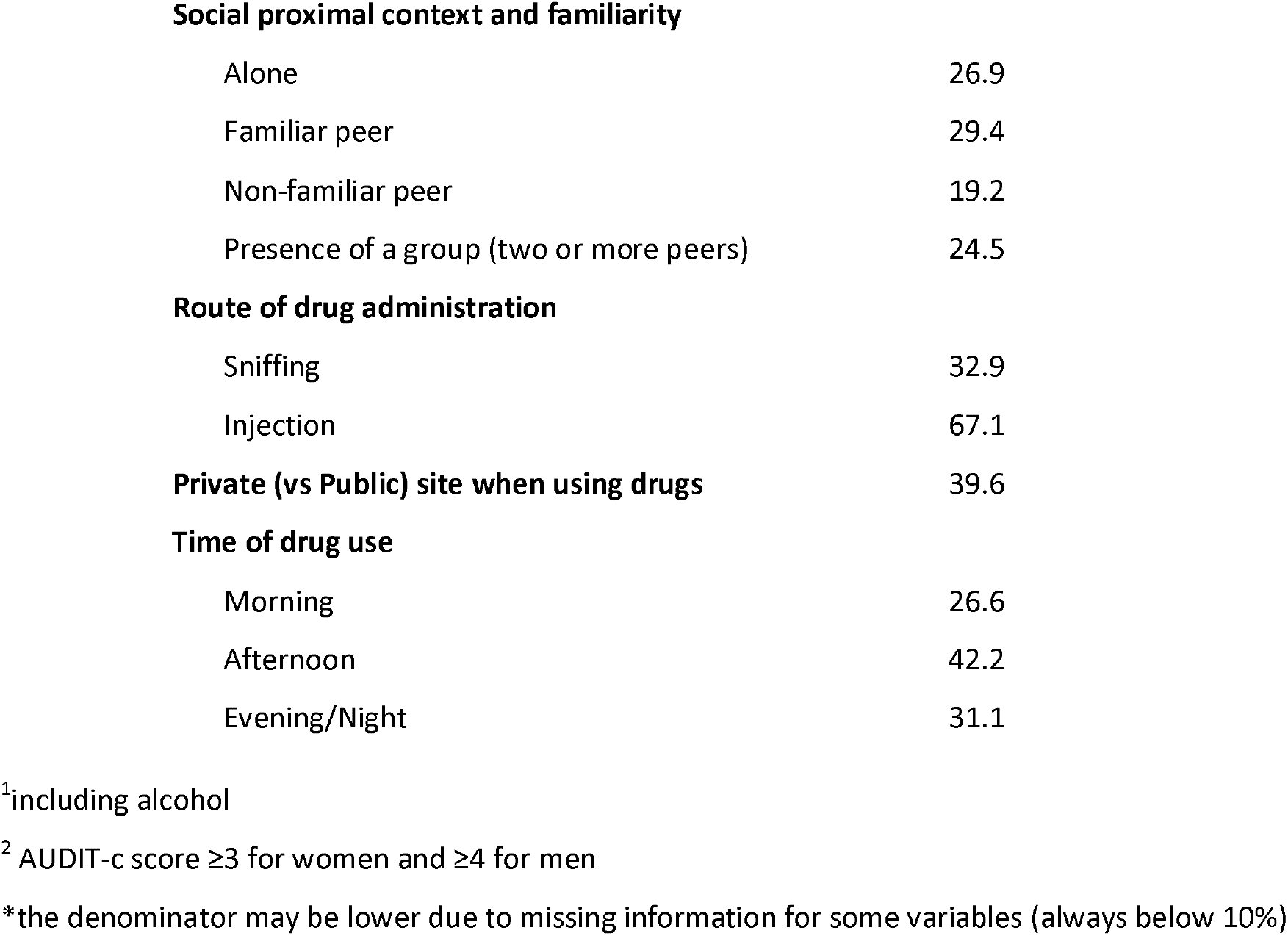
HUMANS- Characteristics of the study sample (N=77) and drug use at any episode reported by participant (N=246)

|  | **Number of**<br>**participants**<br>**N=77**<br>**% or**<br>**median [IQR]** |
| --- | --- |
| **Male gender** | 83.1 |
| **Age – *years*** | 41 [34; 47] |
| **Daily stimulant use** | 36.4 |
| **Age at first drug use** | 17 [15 ; 22] |
| **>20 cigarettes per day** | 44.7 |
| **Hazardous alcohol use<sup>2</sup>** | 48.7 |
| **Stable housing** | 41.6 |
| **Financial problems** | 66.2 |
| **High school certificate** | 32.0 |
|  | **Number**<br>**of**<br>**episodes**<br>**N=246\***<br>**%** |
| **Social proximal context** |  |
| Alone | 26.8 |
| Presence of one peer | 48.8 |
| Presence of a group (two or more peers) | 24.4 |
| **Social proximal context and familiarity** |  |
| Alone | 26.9 |
| Familiar peer | 29.4 |
| Non-familiar peer | 19.2 |
| Presence of a group (two or more peers) | 24.5 |
| **Route of drug administration** |  |
| Sniffing | 32.9 |
| Injection | 67.1 |
| **Time of drug use** |  |
| Morning | 26.6 |
| Afternoon | 42.2 |
| Evening/Night | 31.1 |

<sup>1</sup>including alcohol<sup>2</sup> AUDIT-c score $\geq 3$ for women and $\geq 4$ for men
\*the denominator may be lower due to missing information for some variables (always below 10%)

**Table S2.A.**
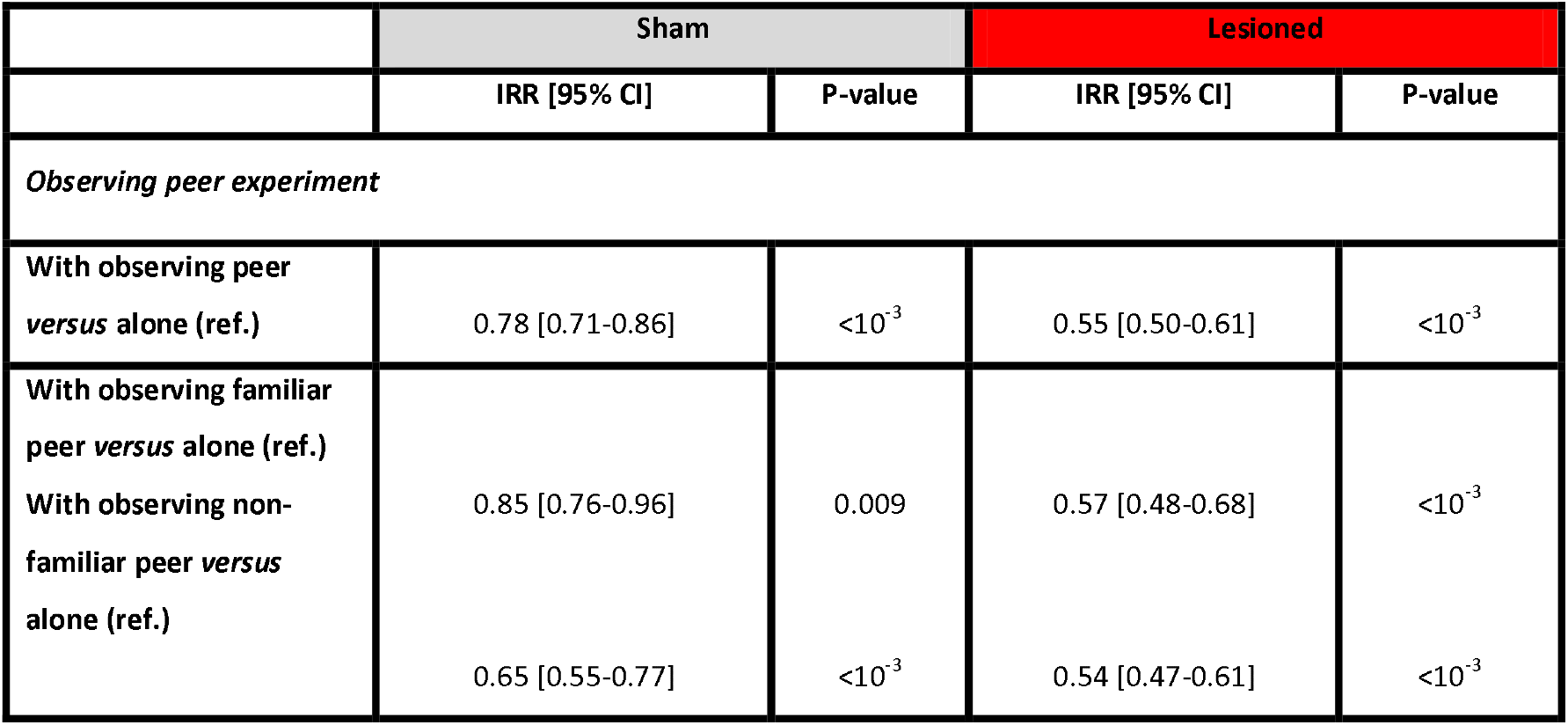
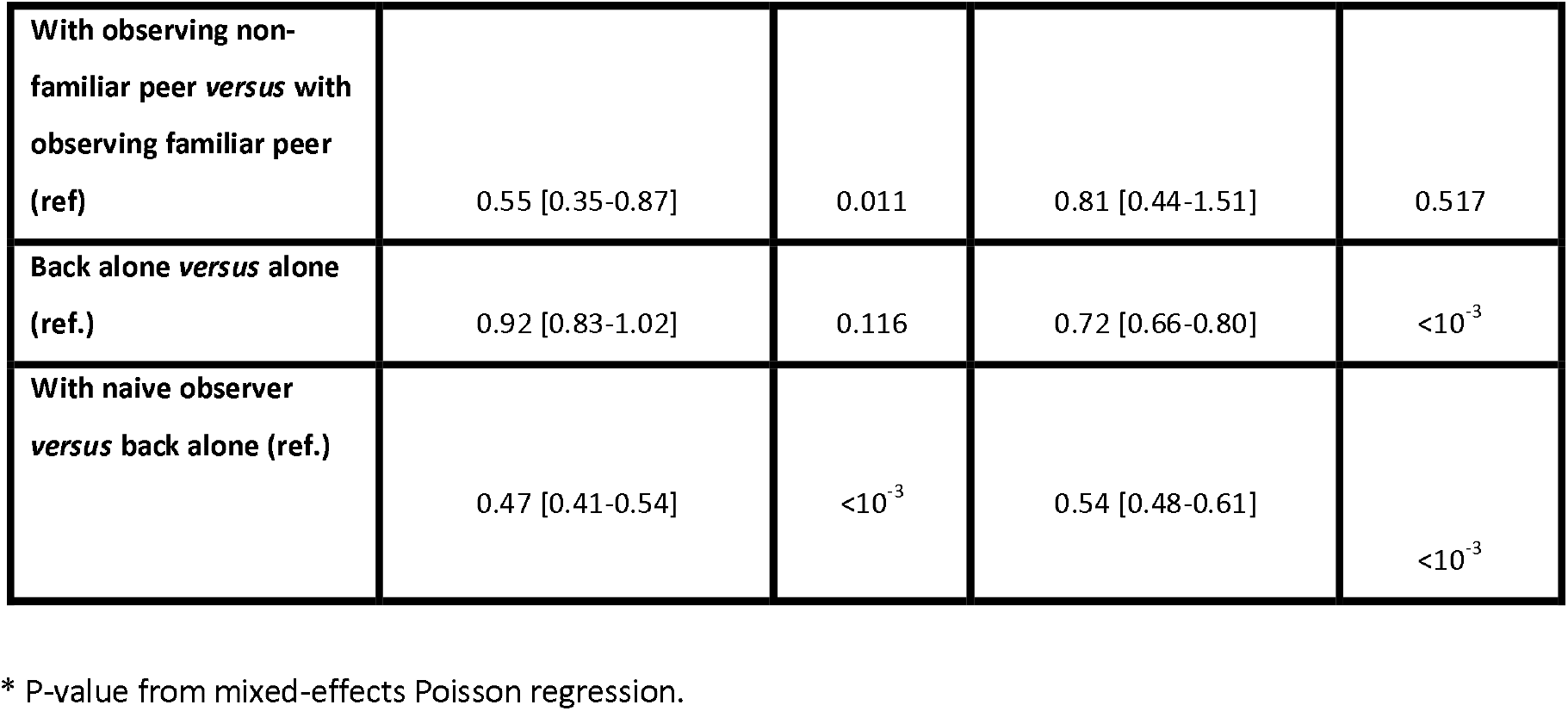

**Table S2.B.**
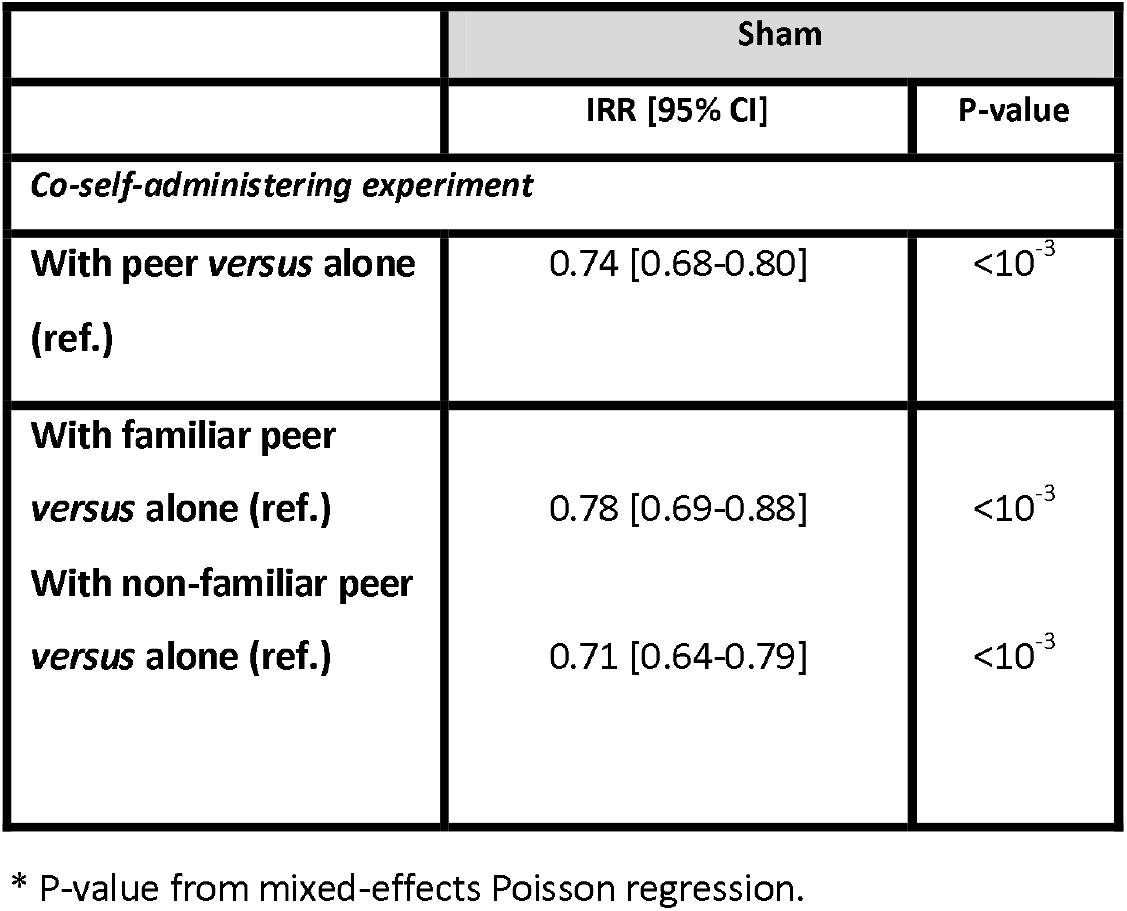

**Fig S1:**
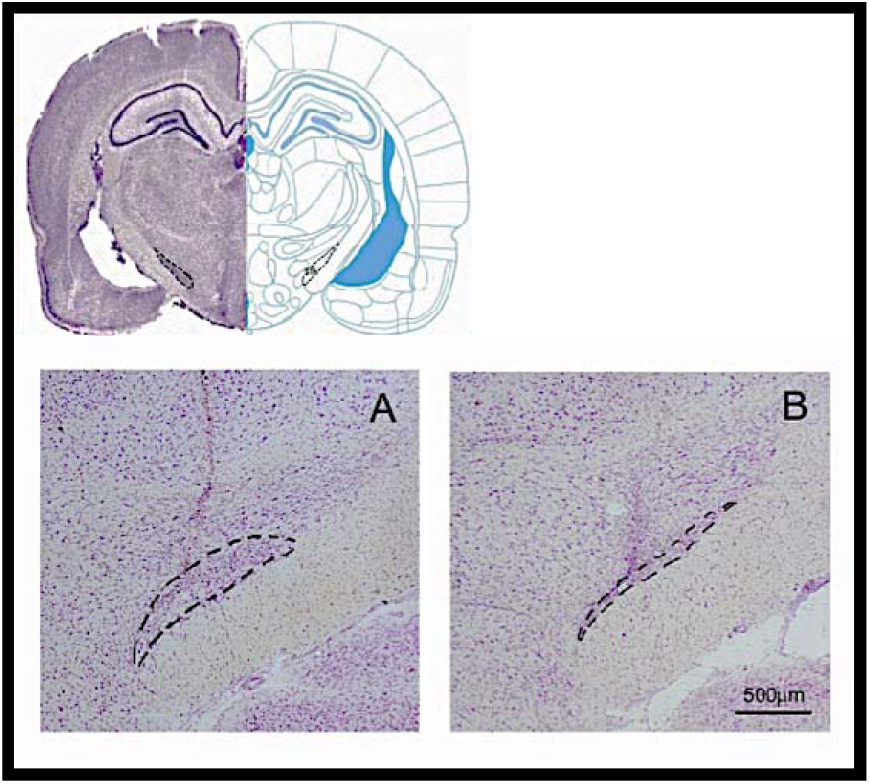
**Up:** Histological brain slice stained with Cresyl violet (left) showing STN position (delineated by dotted lines) and its representation in Paxinos and Watson stereotaxic atlas at the level of the targeted coordinates (right). **Bottom:** Photograph of a brain slice stained with cresyl violet and zoomed at the STN (delineated by dotted lines) level **A**: a sham representative animal. **B**: a STN-lesion representative animal. The STN has shrinked and exhibits gliosis reaction.

